# Eigenmodes of the brain: revisiting connectomics and geometry

**DOI:** 10.1101/2024.04.16.589843

**Authors:** L. Sina Mansour, Hamid Behjat, Dimitri Van De Ville, Robert E. Smith, B.T. Thomas Yeo, Andrew Zalesky

## Abstract

Eigenmodes can be derived from various structural brain properties, including cortical surface geometry^1^ and interareal axonal connections comprising an organism’s connectome^2^. Pang and colleagues map geometric and connectome eigenmodes to spatial patterns of human brain activity, assessing whether brain connectivity or geometry provide greater explanatory power of brain function^3^. The authors find that geometric eigenmodes are superior predictors of cortical activity compared to connectome eigenmodes. They conclude that this supports the predictions of neural field theory (NFT)^4^, in that “brain activity is best represented in terms of eigenmodes derived directly from the shape of the cortex, thus emphasizing a fundamental role of geometry in constraining dynamics”. The experimental comparisons favoring geometric eigenmodes over connectome eigenmodes, in conjunction with specific statements regarding the relative efficacy of geometry in representing brain activity, have been widely interpreted to mean that geometry imposes stronger constraints on cortical dynamics than connectivity^5–9^. Here, we reconsider the comparative experimental evidence focusing on the impact of connectome mapping methodology. Utilizing established methods to mitigate connectome construction limitations, we map new connectomes for the same dataset, finding that eigenmodes derived from these connectomes reach comparable accuracy in explaining brain activity to that of geometric eigenmodes. We conclude that the evidence presented to support the comparative proposition that “eigenmodes derived from brain geometry represent a more fundamental anatomical constraint on dynamics than the connectome” may require reconsideration in light of our findings. Pang and colleagues present compelling evidence for the important role of geometric constraints on brain function, but their findings should not be interpreted to mean that geometry has superior explanatory power over the connectome.

## High-resolution connectome mapping

Pang and colleagues analyzed connectomes mapped at very high resolution (∼32k vertices/nodes per hemisphere). High-resolution connectome mapping is challenging and susceptible to biases and inaccuracies^10,11^. For example, assigning streamline endpoints to vertices of the cortical surface mesh implies accurate identification of endpoint locations with a precision of the inter-vertex distance. Accumulation of integration errors during streamline propagation can exceed this tolerance, leading to unreliable connectivity estimates. While methods are available to alleviate this source of inaccuracy^12^, they do not appear to have been used by Pang and colleagues. Another consideration is gyral bias^13^ –the tendency for streamlines to preferentially terminate at gyral crowns rather than sulcal fundi, leading to biased connectivity estimates. Gyral bias is visibly prominent for the connectomes used by Pang and colleagues (Fig.2b of Pang et al.^3^, also see Supplementary Fig. S1).

Additionally, the authors analyzed binarized connectomes, where continuous connectivity strength information was converted to binary values. While binarization simplifies the connectome, it may result in information loss and reduced robustness to tractography inaccuracies. Given these observations, we sought to evaluate the impact of addressing these issues on the “comparatively poor performance of connectome eigenmodes”.

We mapped new connectomes for the same individuals analyzed by Pang and colleagues, using established methods to alleviate gyral bias as well as to improve streamline assignments and tractography accuracy. Our pipeline included: i) combined intensity normalization and bias field correction of the diffusion MRI data^14,15^; ii) anatomically constrained tractography with streamlines seeded from the white-gray matter boundary using tissue-type segmentation to improve tractography accuracy^16^; iii) gyral bias^13^ reduction via these changes to the tractography pipeline (steps i and ii) relative to the pipeline used by Pang and colleagues, or regression of streamline counts against cortical curvature during connectome postprocessing; iv) retainment of connectome weights to preserve the broad range of interareal connection strengths^17^, rather than reducing to an oversimplified binary connectivity representation^18^; and, v) connectome spatial smoothing to account for imprecision in streamline endpoint determination^12^. We mapped connectome eigenmodes from weighted connectomes pruned to a connection density of 10% and evaluated the impact of alternative densities (see Supplementary Material: Factors influencing reconstruction accuracy of connectome eigenmodes). The above methods are established and commonly used to reconstruct connectomes for purposes other than computing eigenmodes^10–18^.

## Connectome and geometric eigenmodes can explain brain activity equally well

Our connectome eigenmodes explained brain activity with substantially higher accuracy than those used by Pang and colleagues for both resting-state (AUC; our connectomes: 74.8% [74.4% to 75.2%], Pang: 65.7% [65.1% to 66.2%]) and task conditions (AUC; our connectomes: 84.0% [82.9% to 85.0%], Pang: 78.2% [76.7% to 79.7]). Critically, our connectome eigenmodes performed equally as well as geometric eigenmodes for both resting-state (AUC; our connectomes: 74.8% [74.4% to 75.2%], geometry: 75.5% [75.1% to 75.9%]) and task conditions (AUC; our connectomes: 84.0% [82.9% to 85.0%], geometry: 83.1% [82.1% to 84.1%]). This suggests that the “comparatively poor performance of the connectome eigenmodes” used by Pang and colleagues may be a consequence of unaddressed challenges in mapping high-resolution connectomes (Fig. 1, see Supplementary Material for detail).

**Fig. 1.**
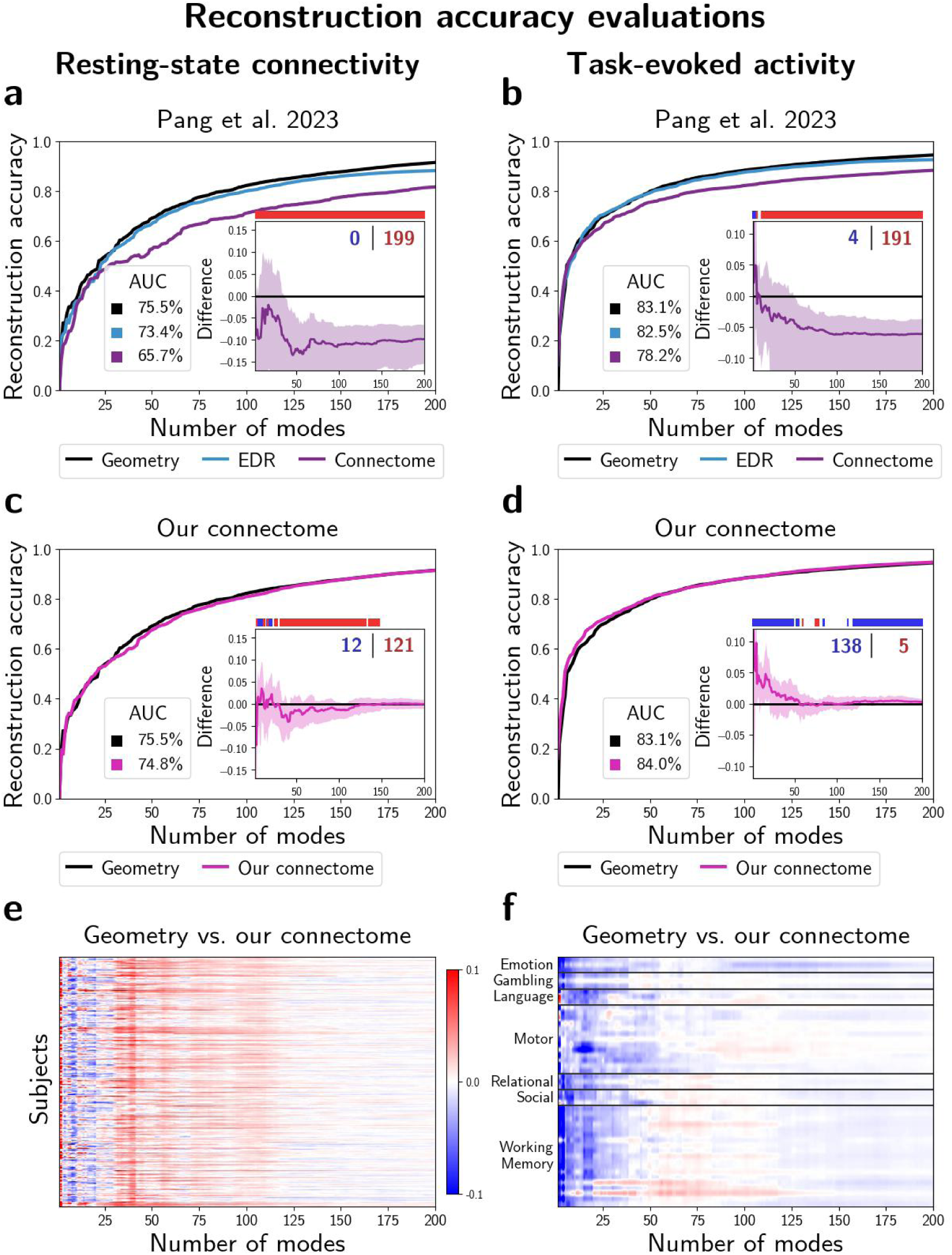
Connectome and geometric eigenmodes explain resting-state and task-based brain activity equally well. (a, b) Replication of results by Pang and colleagues. Plots show accuracy in explaining resting-state (a) and task-evoked activity (b) as a function of the number of eigenmodes. Area under curve (AUC) is shown to provide a summary of reconstruction accuracy. Inset shows the difference in reconstruction accuracy between geometric and connectome eigenmodes. Shading indicates 95% confidence intervals. A nonparametric paired test was used to assess statistical significance; instances where the connectome (blue) or geometry (red) provide a significantly higher reconstruction accuracy (FDR corrected) are marked above the inset and the total counts are reported in the top right corner. (c, d) Same as (a,b), but for our connectome eigenmodes. (e, f) Heatmaps show differences in reconstruction accuracy between geometric and our connectome eigenmodes across individual subjects and task contrasts. Red indicates superior explanatory power of geometric eigenmodes; blue indicates superior explanatory power of connectome eigenmodes.

Reconstruction accuracy differences between our connectome eigenmodes and geometric eigenmodes were modest. These differences were greatest for low-frequency eigenmodes, but converged to zero at higher frequencies, never exceeding a difference of 1% after inclusion of 150 eigenmodes or more. Supplementary analyses were conducted to determine the relative impact on reconstruction accuracy of each connectome mapping step. Using weighted connectomes and controlling for gyral bias were the most important steps in achieving accurate connectome eigenmode reconstructions (supplementary Figs. S2-S10).

## Comparison of connectome and geometric eigenmodes

The spatial profiles of our connectome eigenmodes were similar to that of the geometric eigenmodes, whereas weaker spatial similarity was evident between geometric eigenmodes and the connectome eigenmodes used by Pang and colleagues (see Fig. 2 and supplementary Fig. S11). Our results indicate a greater degree of overlap between brain geometry and our connectome eigenmodes. Shared information between geometry and connectivity may be a contributing factor to the high reconstruction accuracy of both bases. We also carried out supplementary analyses to evaluate the influence of connection length on the explanatory power of connectome eigenmodes (supplementary Figs. S12, S13). Crucially, despite the similarity between geometry and connectome eigenmodes, partial correlation evaluations indicate that both long and short anatomical connections enable connectome eigenmodes to capture sources of spatial variance in brain activity that elude geometric counterparts.

**Fig. 2.**
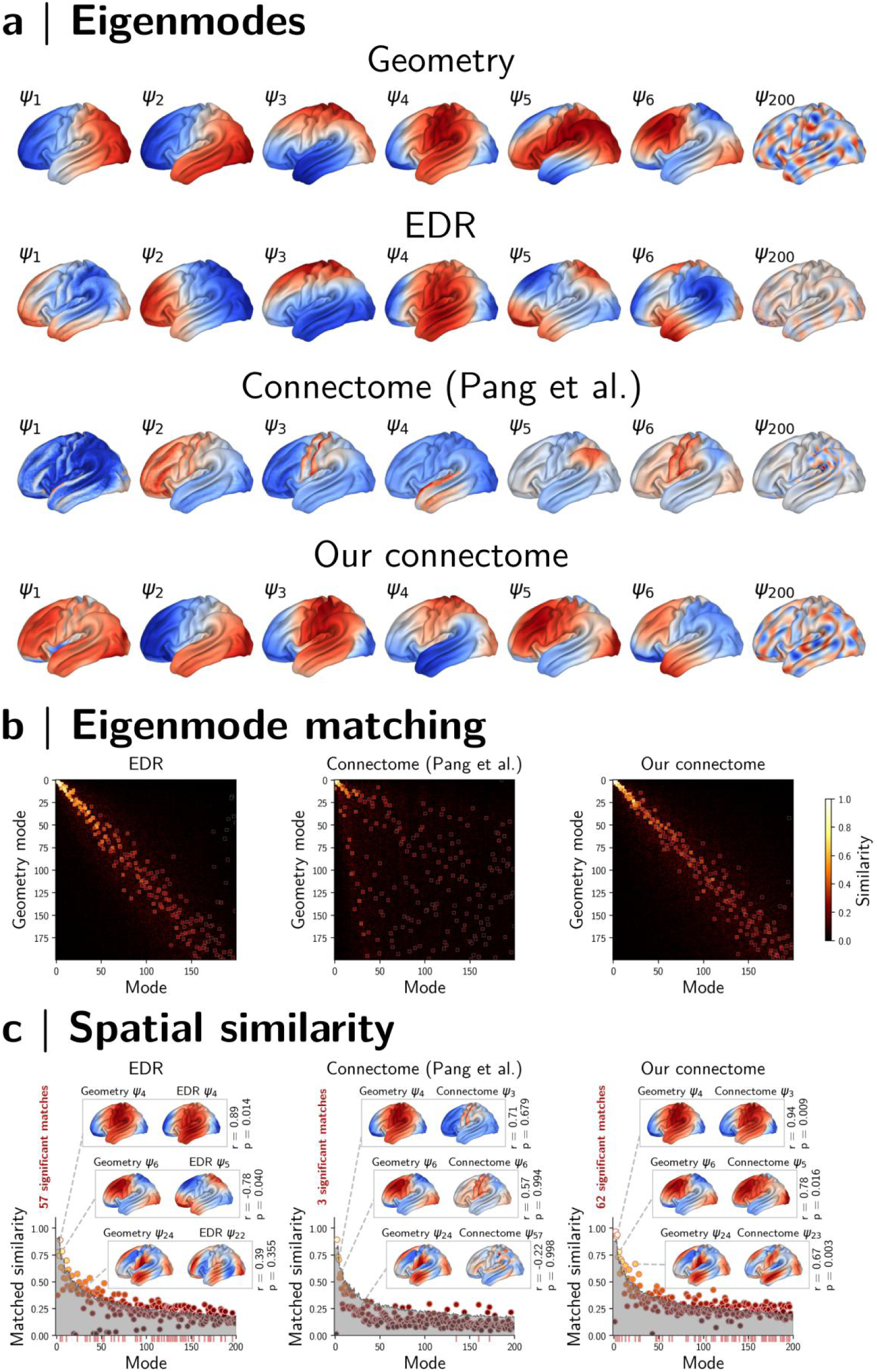
Spatial comparison of geometric and connectome eigenmodes. (a) Cortical surface rendering of geometric (first row), exponential distance rule (EDR, second row) and connectome (third row) eigenmodes mapped by Pang and colleagues. Final row shows eigenmodes derived from our connectomes. (b) The connectivity-based eigenmodes (EDR and connectome) were compared to the geometric eigenmodes. Heatmaps show the spatial similarity across eigenmode pairs, quantified by the magnitude of the Pearson’s correlation. Similar eigenmodes were matched by maximum weighted matching. The squares denoting matched eigenmode pairs are magnified in the heatmaps. (c) A permutation-based nonparametric test was used to evaluate the statistical significance of observed similarities compared to null signals of similar spatial frequency. The gray shade indicates the 95% confidence interval as per the permutaion test. Red ticks along x axis indicate matched pairs with significant similarities (FDR corrected p<0.05) and the total number of significant matches is depicted in red above y axis. The insets present three exemplar pairs of matched cortical eigenmodes (matched to geometric modes 4, 6, and 24).

## Concluding remarks

Pang and colleagues assert that “structural eigenmodes derived solely from the brain’s geometry provide a more compact, accurate and parsimonious representation of its macroscale activity than alternative connectome-based models”. We mapped connectomes for the same individuals using sophisticated techniques^10–18^ and found that connectome eigenmodes are as effective as geometric eigenmodes in explaining resting-state and task-evoked brain activity. The comparatively poor performance of the connectomes used by Pang and colleagues may therefore be attributed to unaddressed concerns in connectome reconstruction. Connectome eigenmodes provided marginally higher reconstruction accuracy than geometry at low frequencies (first 10-30 eigenmodes), underscoring the potential significance of structural connections in shaping large-scale functional hierarchies. Nonetheless, we view these modest differences as insufficient grounds to substantiate conclusions on superiority.

We found similarities in the cortical spatial profiles of connectome and geometric eigenmodes, suggesting a tight relationship between cortical geometry and cortico-cortical connectivity. White-matter connections reflect geometric characteristics, wherein regions in closer spatial proximity exhibit stronger connectedness^14^ (see Fig. S3). Conversely, the brain’s geometric features may be sculpted by interareal connectivity; it is hypothesized that the forces generated by axonal elongations gradually shape cortical folding patterns^15^. The interplay between brain geometry and connectivity remains a fertile area for future exploration. We found that short-range connections were important to connectome explanatory power at higer frequencies; short connections are likely to conform to cortical surface geometry, and therefore disambiguating the effects of connectivity and geometry across short ranges will be challenging.

The work of Pang and colleagues has been construed as pitting the geometric and connectome eigenmode bases against one another^5–8^, even if this was not the authors’ intention^9^. While geometric eigenmodes may be more parsimonious than their structural connectome counterparts, our results demonstrate that specific comments predicated on an inferior explanatory capacity of connectome eigenmodes should be tempered to avoid continued misinterpretation. Our results also highlight challenges of connectome mapping and the importance of utilizing state-of-the-art connectome reconstruction techniques for broad conclusions regarding brain structure and function to be robust.

## Code and data availability

We have made all our code and supplementary data publicly available to facilitate replication of these complementary analyses. The supplementary code and data to this commentary is available from the following repository: https://github.com/sina-mansour/brain_eigenmodes

## Acknowledgements

The analysis for this work was supported by Spartan High-Performance Computing infrastructure, and dedicated computing solutions provided by the Research Computing Services at the University of Melbourne. We gratefully acknowledge the invaluable contribution of several open-source software packages that significantly facilitated our data analysis and interpretation. Specifically, the tractography pipeline was developed using MRtrix3 software^21^. The analytical pipeline made use of several open source python packages including Nibabel^22^, NumPy, and SciPy^23^. The connectome spatial smoothing package^24^ was used for generation and spatial smoothing of high-resolution connectomes. Reproducible brain visualizations were generated by Cerebro brain viewer^25^. RS is a fellow of the National Imaging Facility, a National Collaborative Research Infrastructure Strategy (NCRIS) capability, at the Florey Institute of Neuroscience and Mental Health.

## Supplementary Material

### Eigenmode reconstruction coefficients

We estimated geometric eigenmode coefficients using the linear regression model outlined in the supporting scripts openly provided by Pang and colleagues. Linear regression was used because the geometric eigenmodes provided by the authors were not orthonormal (0<|r|<0.1). In contrast, graph Laplacian-based eigenmodes (e.g., EDR and connectome) were orthonormal (r=0) and thus the dot product was used to estimate eigenmode coefficients, as described in equations 4,5 of Pang et al. (2023). We note that the coefficients from linear regression (without an intercept) and the dot product are equivalent if the eigenvectors are orthonormal.

### Factors influencing reconstruction accuracy of connectome eigenmodes

We mapped connectomes for the same individuals analyzed by Pang and colleagues to generate connectome eigenmodes and reconstruct individual resting-state functional connectivity matrices and task-evoked activity maps (for 47 different task contrasts). As discussed above, we implemented established methods to alleviate several biases in the connectome mapping pipeline^5–13^. Subsequent sections will detail these procedures and evaluate their specific influence on reconstruction accuracy.

### Connectome density

Binarizing connectomes at lower densities, particularly when utilizing group-average connection strength as the threshold criterion, may result in the unintended removal of genuine structural connections. Notably, due to the exponential decay in the prevalence of connections as a function of streamline length^12,14^, a stringent threshold (low density) is likely to disproportionately eliminate long-range connections^26,27^. In line with evaluations reported by Pang and colleagues^3^, our findings (Fig. S2.b) verify that increasing the binarization density (from 0.1% to 1%) yields improvements in connectome eigenmode reconstruction accuracy for rest (AUC: from 65.7% to 71.7%) and task conditions (AUC: from 78.2% to 81.6%).

### Alleviating gyral bias

Upon inspection, the surface projections of connectome eigenmodes manifested noticeable gyral bias (Fig. S1, also visible in Fig.1 of the original article). Gyral bias is a known issue in tractography whereby regions located on the gyral ridges receive a proportionally higher number of streamlines relative to those in the gyral wall and sulci^8^. After thresholding, gyral bias may lead to disproportionate removal of sulcal connections, and hence underrepresenting sulcal connectivity. Previous studies have proposed different strategies to mitigate this bias^28,29^. Here, we used two alternative approaches to control for this bias and assessed their influence on reconstruction accuracy. First, we performed a linear adjustment to regress cortical curvature from connectivity strength. This correction step was performed before density thresholding to mitigate the risk of excessive removal of sulcal connections. As presented in Fig. S2.c, this correction for binarized connectomes results in a further increase in reconstruction accuracy of connectome eigenmodes for both rest (AUC: from 71.7% to 73.1%) and task (AUC: from 81.6% to 82.9%) conditions. We also considered an alternative tractography pipeline to reduce the gyral bias (detailed in the ensuing section).

### Tractography pipeline

To investigate whether the underperformance of connectome eigenmodes was attributable to gyral and other tractography biases, we employed an alternative tractography pipeline to minimize the impact of such biases. Streamline tractography was conducted with MRtrix3^21^ and adopted previously used steps detailed elsewhere^30^. In contrast to the pipeline used by Pang and colleagues, we incorporated combined intensity normalization and bias field correction of the diffusion-weighted imaging data^9,10^ and seeded streamlines from the white-gray matter boundary computed from 5-tissue-type segmentation to perform anatomically constrained tractography (ACT)^11^. As illustrated in Fig. S1, these steps substantially reduced gyral bias. As further corroborated in Fig. S2.d, when gyral and other tractography biases are alleviated, connectomic eigenmodes demonstrate enhanced reconstruction performance, obviating the need for explicit curvature adjustments as discussed in the preceding section. Specifically, to evaluate the incremental benefit of the tractography changes, a group average weighted connectivity matrix was constructed from the new tractograms, followed by binarization using a 1% density threshold (without regression of gyral bias). Resulting binary connectomes yielded eigenmodes with increased reconstruction accuracy across both rest (AUC: from 71.7% to 74.2%) and task (AUC: from 81.6% to 83.1%) conditions. This verifies that mitigating gyral bias, whether via linear confound regression (Fig. S2.c) or tractography procedures (Fig. S2.d) improves reconstruction accuracy.

### Smoothed weighted connectivity

Pang and colleagues used binarized matrices to estimate the connectome eigenmodes^3^. Binarization can obscure meaningful variations in connectivity strengths. Studies using tractography and tract tracing techniques have previously indicated that interareal connection strengths vary by multiple orders of magnitude^12,14^. This is particularly evident in the high-resolution connectomes discussed here (Fig. S3); such that binarized high-resolution connectomes may fail to include long-range structural connections due to their comparatively lower strengths. Established connectome mapping guidelines assert that binarization may oversimplify the connectivity matrix and recommend the adoption of weighted connectomes^13^. We therefore computed Laplacian eigenmodes using weighted connectomes. First, connectome spatial smoothing (8mm FWHM) was applied to enhance the interindividual reliability of high-resolution connectomes and reduce streamline endpoint location innacuracies^6,31^. Compared to binary connectomes, where connectome density can strongly influence network topology and attributes, removal of the weakest elements from a weighted structural connectome matrix has only minor influence, due to the range of the connection strength distribution^32^. As such, following connectome smoothing (which intrinsically increases connectome density, Fig S3), we used a more lenient threshold to prune the connectomes at 10% density prior to eigenmode estimation.

To construct the adjacency matrix, we integrated this connectivity data with local connections, akin to the authors’ approach^3^. This step is requisite to form a fully connected matrix, which, in turn, is requisite for the eigenmode calculation. While the binary version employed by the authors utilized a logical OR operator, we use the equivalent weighted operation of summation (W*_c_* = W*_connectome_* + *ε_local_* A*_local_*, where W*_c_* denotes the matrix that combines tractography-based connectivity with local vertex adjacency). The local connection weights were multiplied by a small scaler (*ε_local_* = 10^−6^) to ensure that the results were primarily influenced by the connectome weights. The eigenmodes resulting from this final step served as our alternative connectome eigenmodes. Notably, when comparing the gain of this last step against the eigenmodes constructed from binarization of the updated tractography pipeline (previous section), the smooth weighted alternative connectome eigenmodes resulted in relatively modesl performance improvements for both rest (AUC: from 74.2% to 74.8%) and task (AUC: from 83.1% to 84.0%) conditions (see Fig. S2.e).

### Systematic evaluation of parameters

Our analyses indicated that several decisions in structural connectome reconstruction contribute to the observed improvements. To quantify the sensitivity of these improvements to different parameter choices, we conducted a systematic evaluation (see Figs. S4 to S10). Given the large number of possible parameter combinations across the connectome mapping pipeline, it is computationally infeasible to explore every permutation (e.g., connectome density, gyral bias correction, tractography pipeline, binary vs. weighted, smoothing strength, global-local combinations). As a result, we systematically examined the impact of changing a single parameter while keeping other parameters constant.

Figs S4, S5, & S6 evaluate the effect of binary connectome density on different connectomes (with or without gyral bias correction). This shows that densities between 0.5% to 1% consistently yield high reconstruction acccuracies when constructing eigenmodes from binary connectomes (regardless of gyral bias correction). Moreover, we also evaluated the effect of density for pruned weighted connectomes (Fig S8); our results indicate that when using weighted connectomes, densities above 0.5% are remarkably better than lower densities. In contrast to binary counterparts, increasing the density of weighted eigenmodes above 1% did not detriment reconstruction accuracy (Fig S8). Figs S7 and S9 examine the impact of performing connectome spatial smoothing. Notably, using connectome smoothing, particularly with wide kernels (6-10mm FWHM) can result in marginal improvements in reconstruction accuracy. Finally, as shown in Fig S10, we evaluated the impact of changing the global-local combination parameter (*ε_local_*) and found that it had negligible impact on reconstruction accuracy.

### Eigenmode similarity comparison

In Fig. 2, the geometric and connectivity-based eigenmodes were spatially compared to assess the degree of similarity between eigenmodes. To this end, for all connectivity-based eigenmodes, a 200×200 similarity matrix was computed that quantified the absolute value of the Pearson’s correlation between the eigenmode basis set with the geometric eigenmodes. This quantified the degree of spatial correspondence between two sets of eigenmode bases. Next, a maximum weighted matching (via linear sum assignment using a modified Jonker-Volgenant algorithm with no initialization^33,34^) was used to find optimal matching pairs of eigenmodes such that the total spatial correspondence between eigenmodes were maximized. To evaluate the spatial interdependencies between the matched pairs, a permutation-based non-parametric spin-test was implemented^35^. Specifically, for a total of 10,000 permutations, the geometric eigenmodes were collectively randomly rotated to form a spatial null with similar frequency characteristics. For each permutation, a maximum weighted matching method was similarly used to find the matched pairs. This created a null distribution of matched eigenmode similarities for each mode. Next, p-values were generated for every pair of matched eigenmodes to quantify the likelihood of observing that magnitude of similarity in the null distribution. The resulting p-values per matched eigenmode pair were then FDR corrected to find significant similarities across all tests (Benjamini-Hochberg Procedure). This illustrated that both the EDR eigenmodes (57 significant matches) and our connectome eigenmodes (62 significant matches) contained a greater degree of similarity to geometry eigenmodes than expected by chance alone.

### Subspace similarity comparison

The analyses presented in Fig. 2 quantified the degree of spatial correspondence between matched eigenmode pairs. However, not only can the order of eigenmodes differ between two sets of eigenmodes (e.g., comparing geometric eigenmodes with EDR, Pang et al. 2023 connectome, and our connectome eigenmodes) but also a linear combination of eigenmodes from one design can represent an eigenmode from another design. It is therefore important to compare subspaces spanned by sets of eigenmodes from different designs in addition to comparing individual pairs. To this end, we used the Procrustes transform (PT)^36^, which finds the optimal rotation to match two linear subspaces. This method has been previously used in related work^37^, albeit with a different goal, to quantify the degree of inter-subject variability of eigenmodes of voxel-wise brain graphs. Given two sets of K eigenmodes, PT optimally transforms one set to match the other set.

Specifically, we treated the geometric eigenmodes as the reference set (not transformed) and computed the PT that optimally maps the eigenmode of EDR, Pang’s connectome and our connectome eigenmodes to the reference. This procedure was repeated for different subsets of initial eigenmodes (K from 2 to 200). The cosine similarity between each pair of sets of eigenmodes was computed before and after PT, resulting in two K x K matrices for each K and each pair of designs. Noting that the Frobenius norm of the cosine similarity matrices before and after PT is identical, the norm of the off-diagonal elements after PT provides a measure of residual distance between the two spaces. Normalizing the residual by the total Frobenius norm indicates the proportion of similarity in the subspaces spanned by the pairs of eigenmodes. Particularly, the percentage of the residual before PT retained after PT can be treated as a distance measure. The distance measure before and after PT for different pairs of designs and K can be seen in Fig. S11a,b respectively. These results complement the similarity evaluations presented in Fig. 2; notably, EDR and our connectome eigenmodes have a greater similarity to (smaller distance from) the geometric eigenmodes than connectome eigenmodes used by Pang and colleagues.

### Structural connection lengths

Formation of anatomical connections in the central nervous system is fundamentally constrained by wiring cost^38^. Long-range connections are costly and a trade-off between efficient information transfer and minimal wiring cost influences connectome formation. This posits a potential rationale for why geometric eigenmodes perform equally well in reconstructing spatial patterns of brain activity, particularly at shorter wavelengths: they act as a surrogate for the expected presence of short-range local connections. Short connections are the most common, least costly and are also more likely to conform with cortical surface geometry than long-range connections.

We investigated the potential influence of connection length on eigenmode formation as follows. First, we chose a set of streamline length thresholds as 8, 16, 32, 64mm; the connections contained within the five bins formed by these thresholds are shown for an exemplar vertex in Fig. S12.a. Then, for each threshold, we formed two pruned connectomes: one consisting only of streamlines longer than that threshold, and one consisting only of streamlines shorter than that threshold. Connectome eigenmodes were then computed from these pruned connectomes. This facilitated evaluation of pruned connectomes that exclude either long or short connections as a function of the exclusion threshold.

Where a maximal streamline length is imposed and progressively decreased (Fig. S12.b), reconstruction accuracy remains mostly consistent; at least until that maximal length decreases to 8mm at which point this accuracy declines, except for accuracy with fewer number of modes that are most sensitive to removal of longer connections. Conversely, where a minimal streamline length is imposed and progressively increased (Fig. S12.c), this quickly becomes deleterious to the explanatory power of the connectome eigenmodes, particularly with greater numbers of eigenmodes that ideally capture patterns of higher spatial frequency.

There are two topics of discussion that arise from these results; firstly, given that very short connections are far more prevalent than long connections^39,40^, and that the trajectories of such short length connections are strongly influenced by the local cortical geometry^41^, it is possible that the connectivity information encoded in short wavelength connectome eigenmodes closely mimics that of geometric information.

Secondly, given that inclusion of larger numbers of eigenmodes facilitates mapping of higher spatial frequency components, it is intuitive that exclusion of connections of short length will be deleterious to the explanatory power of the resulting eigenmode bases of greater cardinality. As such, pruning connectomes at minimal lengths of 16mm and 32mm resulted in a plateau in reconstruction accuracy, observed after the initial 50 and 20 eigenmodes, respectively (red pointers in Fig. S13.c).

These results are consistent with the subtle differences in reconstruction accuracy between geometric and connectome eigenmodes. Plots of reconstruction accuracy as a function of number of modes often show that connectomes offer slightly greater explanatory power with a small number of modes whereas with a larger basis set geometry performs equally well. Using a small number of modes of long wavelength, the specificity of structural connectivity estimates may provide contrast that is comparably absent from a purely geometric parametrisation; this is supported by the preservation of explanatory power with a small number of connectome eigenmodes even if short connections are eliminated.

Conversely, by using a larger basis that includes short wavelength eigenmodes, geometry provides comparable explanatory power via an indirect proxy for the local structural connectivity that is biologically responsible for constraining functional activity. The strong prevalence of very short connections, along with the dominance of their contribution toward explanatory power of connectome eigenmodes, contraindicates the conventional use of large minimum length thresholds in tractography.

### Partial correlation evaluations

Evaluations of reconstruction accuracy can fail to discern whether distinct eigenmodes reconstruct similar or disparate sources of information. Given the observed comparable explanatory power of geometric and connectome eigenmodes as well as similarities between the two basis sets, we utilised partial correlations to assess the level of distinction between the two. As reconstruction accuracy was quantified by Pearson’s correlation, partial correlation provides a straightforward extension to measure the incremental contribution of either geometric or connectome basis sets while accounting for the explanatory capacity of the other.

We assessed the incremental value of geometry/connectome eigenmodes across varying number of modes: for any fixed number of modes, we calculated partial correlations between the ground truth and reconstructed data while controlling for the reconstruction achieved by the alternative basis set. Results are presented in Fig. S13.a. Both geometry and connectome eigenmodes can elucidate sources of variance that are independent of the other basis set using the same number of modes. The presence of significant additive benefits for either set of modes suggests that specific sources of spatial variance in functional maps may be more effectively captured by that particular set. Across a wide range of possible numbers of modes, connectome eigenmodes consistently uncover significant sources of variance that elude the geometric counterpart. A similar pattern is also apparent for geometric eigenmodes, except in the reconstruction of task-evoked activity with a small number of modes (<25), where geometric eigenmodes provide no additional benefits to connectome reconstructions. This suggests that the similarity in reconstruction performance between the two sets of eigenmode bases is not due to their equivalence; conversely, each set possesses a significant degree of explanatory capacity that is absent in the other.

We further expanded these assessments to explore the impact of imposing length thresholds on the partial correlation of connectome eigenmodes, controlling for the reconstructions based on geometric eigenmodes. Imposing maximal length thresholds to exclude longer connections negatively impacted the incremental benefit of connectome eigenmodes, particularly when fewer number of modes were considered (see Fig. S13.b). Conversely, imposing minimal length thresholds to exclude shorter connections were most detrimental for the additive benefit of connectome eigenmodes when a higher number of modes were considered (see Fig. S13.c).

This further validates the intuitive expectation that longer connections play a pivotal role in shaping lower frequency modes, while shorter connections contribute to the accurate estimation of higher frequency modes. Particularly noteworthy are the insights from Fig. S13.c, underscoring the distinct contribution of long-range connections to the explanatory power of connectome eigenmodes. For instance, even after the exclusion of connections shorter than 64mm—constituting over 95% of all reconstructed streamlines—the remaining long connections form connectome eigenmodes that significantly capture sources of variance not attainable by geometric modes, particularly evident with fewer than 100 modes considered. This underscores the impact of connections of all lengths in shaping the distinct sources of spatial variance captured by connectome eigenmodes.

**Fig. S1.**
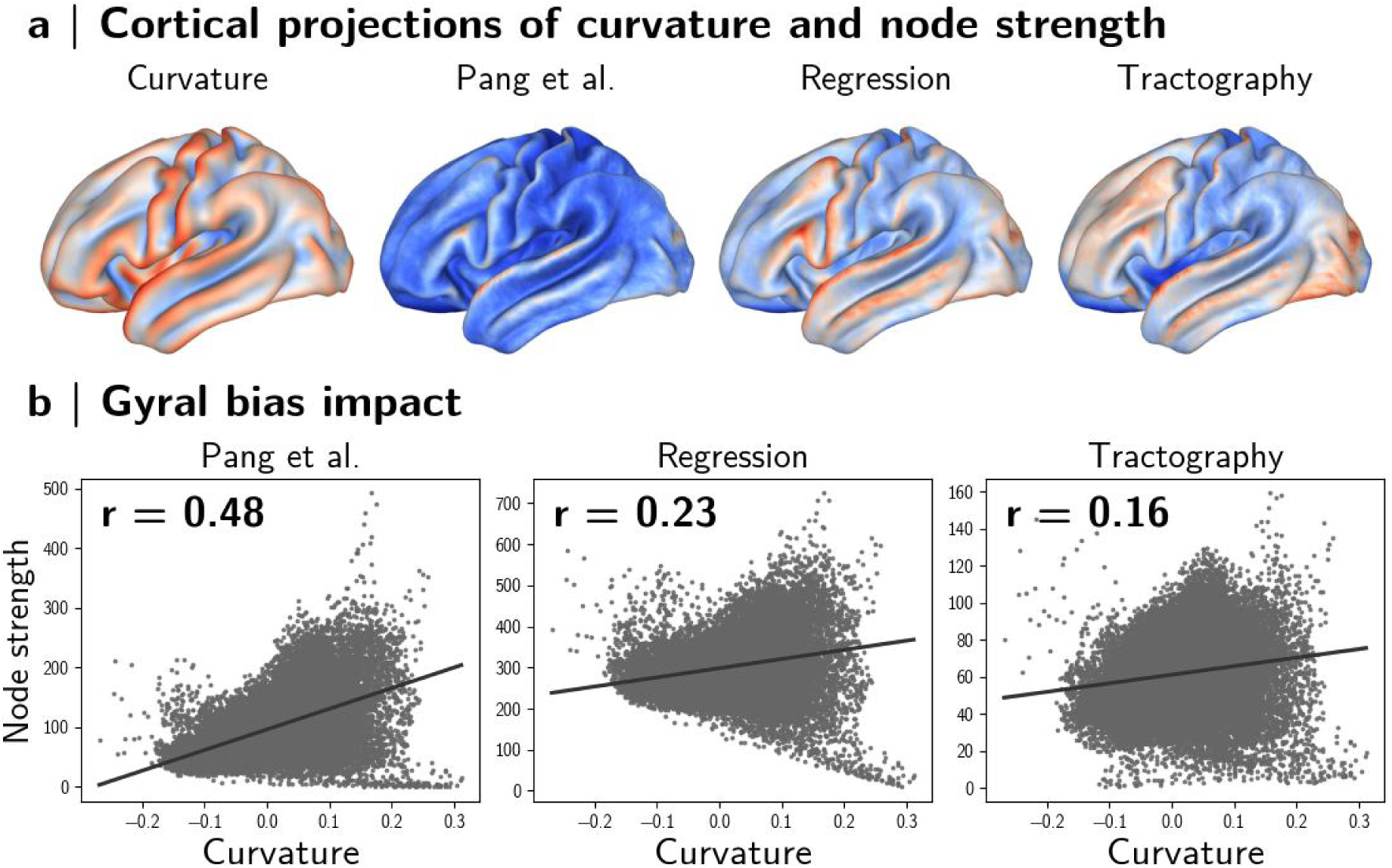
Gyral bias effects visible in connectome strength. (a) Surface projections of cortical curvature (left), connectivity strength, respectively for the original connectome from Pang et al., and connectomes after addressing the gyral bias via regression or tractography. (b)The severity of gyral bias can be quantified by the correlation between curvature and connectivity strength, showing pronounced gyral bias effects in the original connectomes used by Pang and colleagues (left; r = 0.48). In contrast gyral bias is substantially reduced via regression (center; r = 0.23) or tractography (right; r = 0.16).

**Fig. S2.**
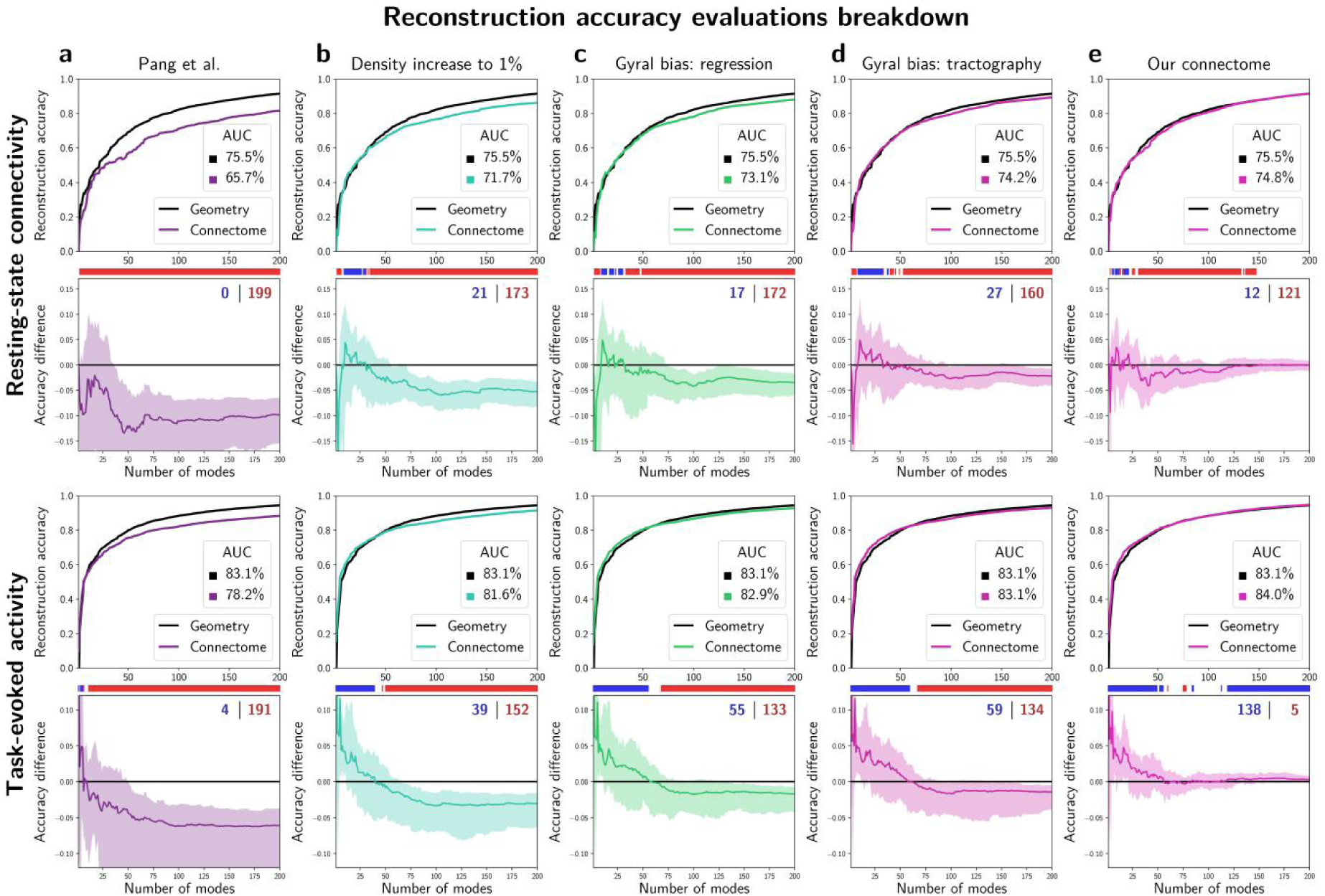
Breakdown of different steps influencing reconstruction accuracy. The format of this figure closely resembles the structure of Figure 2. The first two rows show results for resting state connectivity, and the last two rows show results for task-evoked activity. Both reconstruction accuracy and differences are presented, along with AUC metrics, for ease of comparison. (a) Original results reported by Pang and colleagues. (b-d) Improvement in reconstruction accuracy achieved from eigenmodes of different intermediary connectome construction steps. (b) The impact of connectome density: Increasing the density of binary connectome (from 0.1% to 1%) improves reconstruction accuracy, resulting in a 3-5% increase in AUC. (c) Linear adjustment of edge strength to account for the gyral bias provides additional accuracy improvements, with a 2-3% increase in AUC. (d) Alternatively, similar improvements can also be achieved by mitigating the gyral bias in the tractography pipeline, resulting in a 2-4% increase in AUC. (e) Finally, the use of smoothed weighted connectomes further enhances reconstruction accuracy, leading to a 1% increase in AUC.

**Fig. S3.**
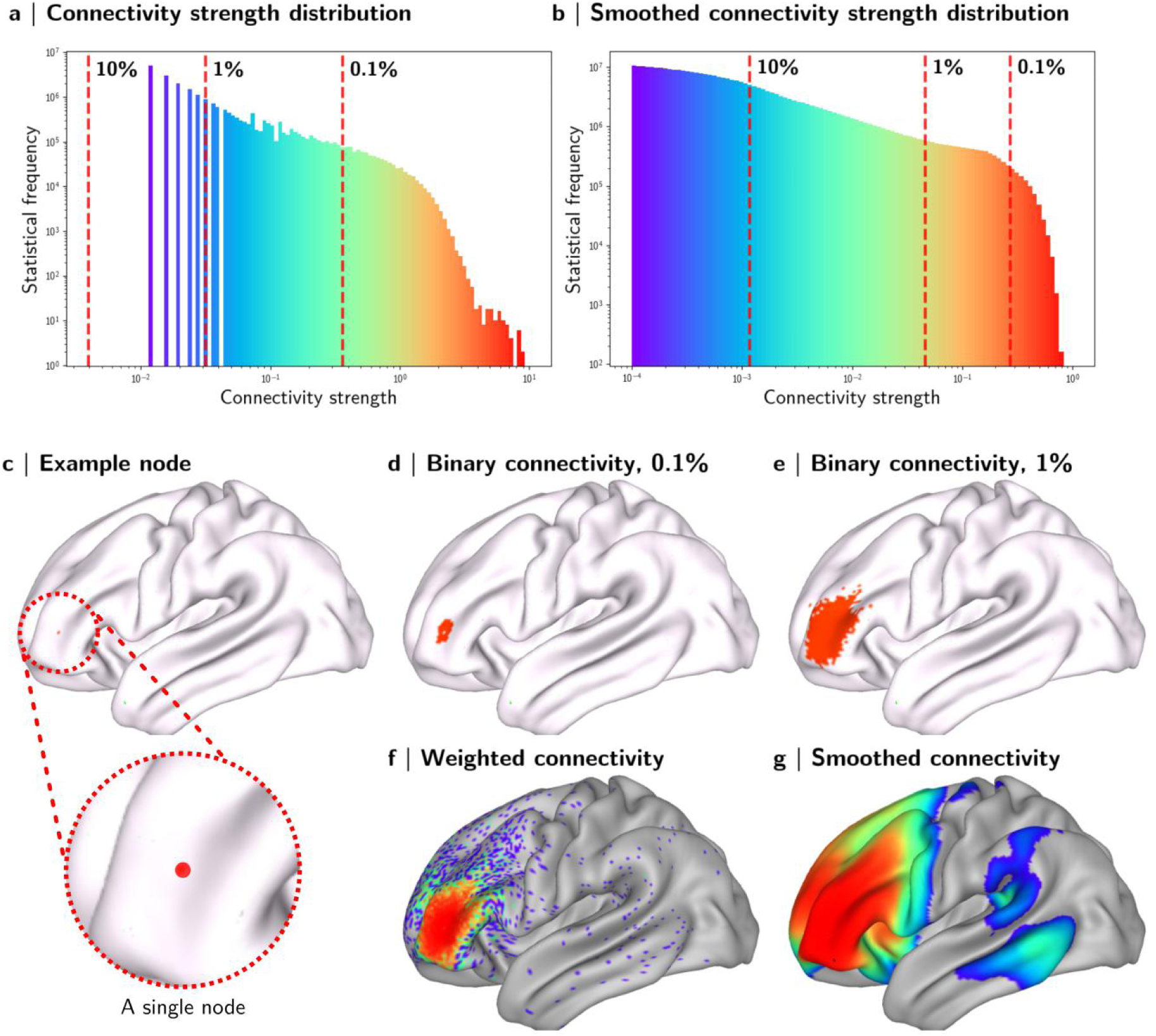
High-resolution connectivity strength spans multiple orders of magnitude. (a,b) Histograms show the distribution of connection strengths in a high-resolution connectome (a) before and (b) after connectome spatial smoothing. Plot axes are log-transformed to highlight the presence of connection strengths spanning multiple orders of magnitude. (c-g) Cortical projections of high-resolution structural connectivity from an exemplary node/vertex situated on the left frontal cortex (c). Binary connectomes simplify the connections of a vertex to a binary mask; this is shown for connectomes binarized at two different density thresholds of 0.1% and 1% (d,e). In contrast, weighted connectomes capture the diverse range of connections with different strength magnitudes which is visible (f) before and (g) after connectome spatial smoothing. This figure also shows the increase in connection density resulting from spatial smoothing. Notably, smoothing improves reconstruction of known long-range connections to the exemplary node, i.e. the superior longitudinal fasciculus and the arcuate fasciculus.

**Fig. S4.**
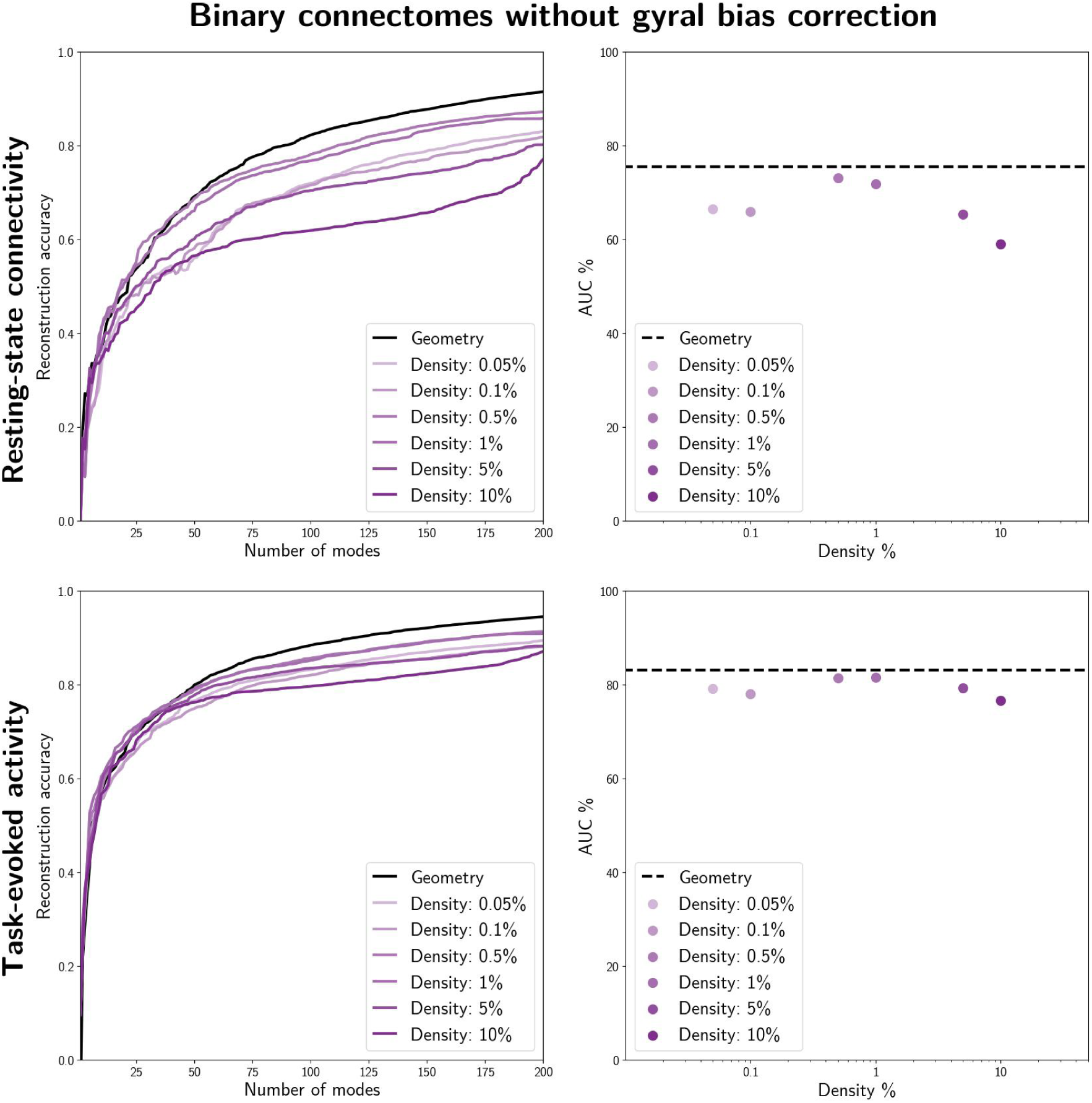
Assessing the impact of density on reconstruction accuracy of binary connectomes without gyral bias correction. Here, the reconstruction task was systematically repeated for varying connectome density levels (0.05% to 10%) while keeping other pipeline parameters constant; specifically, this test utilized the connectomes from Pang et al. (2023) without performing any gyral bias correction, and a binary connectome was constructed based on density thresholds. Line plots (left) depict the reconstruction accuracy as a function of the number of modes for both resting-state connectivity (top) and task-evoked activity (bottom) reconstruction. Scatter plots (right) display the corresponding summary AUC measures. Connectome eigenmode performance is relatively higher within the 0.5% to 1% density range compared to both higher and lower densities.

**Fig. S5.**
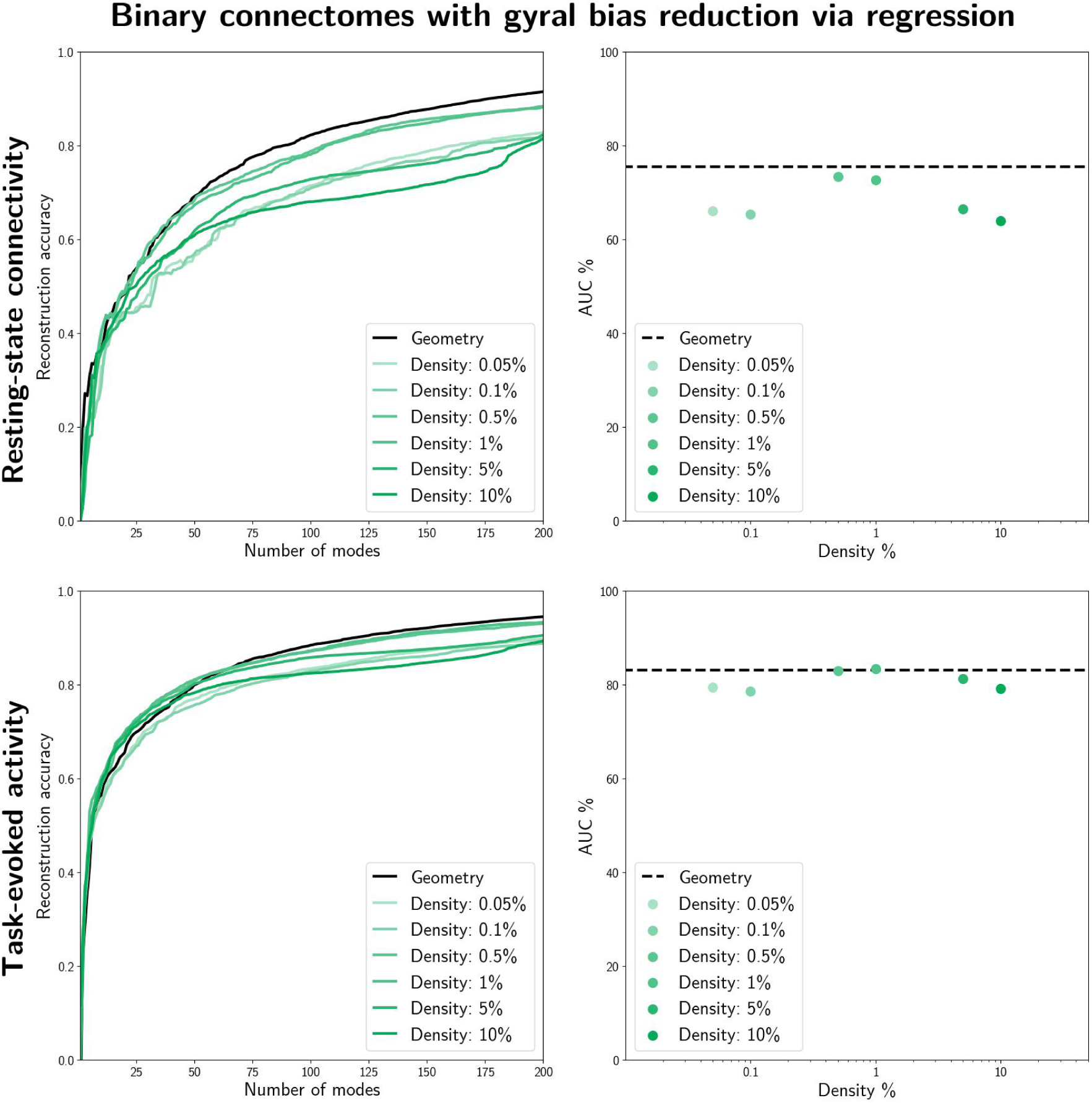
Assessing the impact of density on reconstruction accuracy of binary connectomes after gyral bias reduction via regression. Here, the reconstruction task was systematically repeated for varying connectome density levels (0.05% to 10%) while keeping other pipeline parameters constant; namely, we utilized the connectomes from Pang et al. (2023), applied gyral bias regression, and constructed a binary connectome based on several density thresholds. Line plots (left) depict the reconstruction accuracy as a function of the number of modes for both resting-state connectivity (top) and task-evoked activity (bottom) reconstruction. Scatter plots (right) display the corresponding summary AUC measures. Notably, performance is higher within the 0.5% to 1% density range compared to both higher and lower densities.

**Fig. S6.**
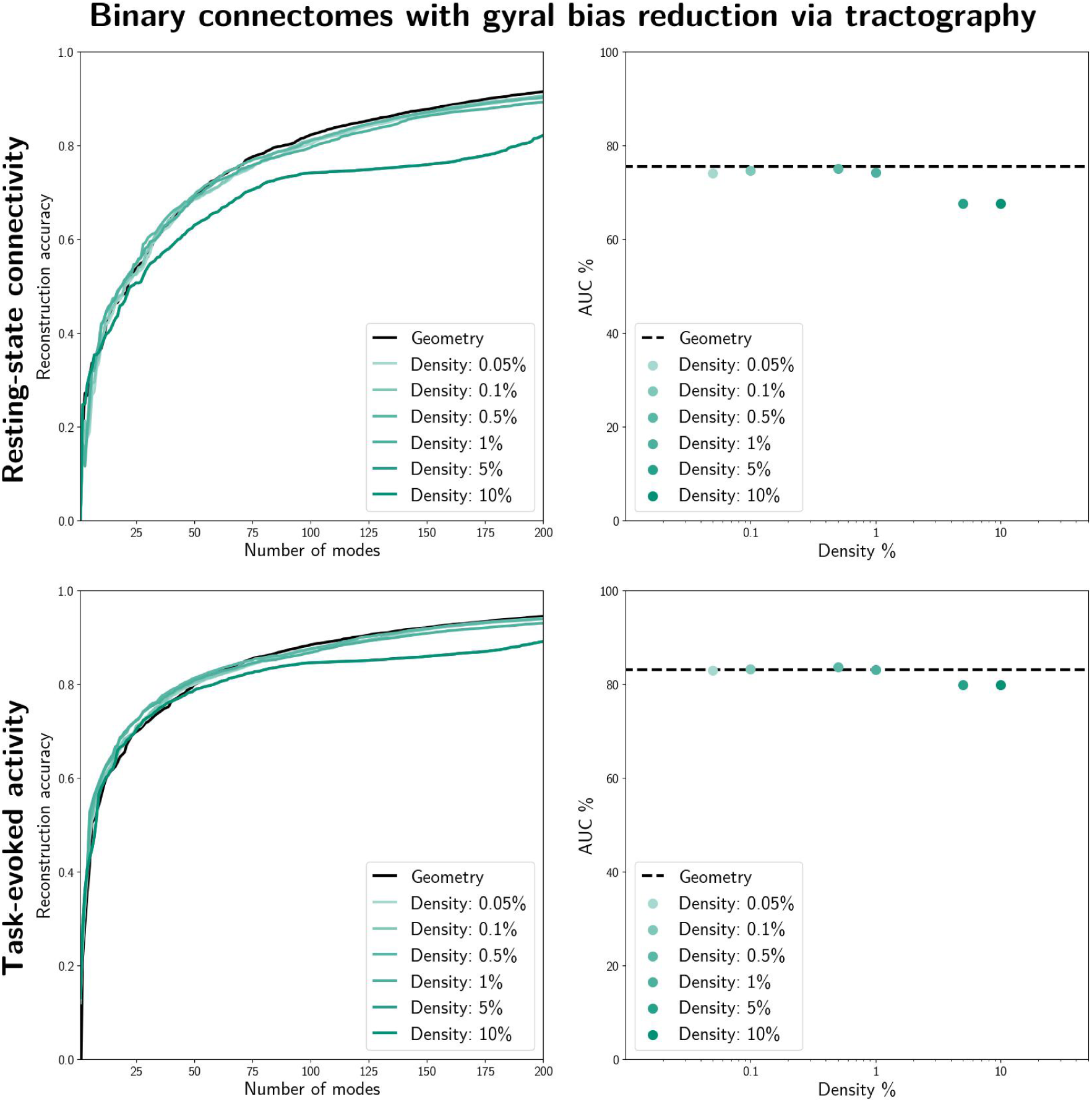
Assessing the impact of density on reconstruction accuracy of binary connectomes after gyral bias reduction via tractography. Here, the reconstruction task was systematically repeated for varying connectome density levels (0.05% to 10%) while keeping other pipeline parameters constant; namely, we utilized the connectomes from our tractography pipeline that better mitigated the gyral bias, and constructed a binary connectome at several density thresholds. Line plots (left) depict the reconstruction accuracy as a function of the number of modes for both resting-state connectivity (top) and task-evoked activity (bottom) reconstruction. Scatter plots (right) display the corresponding summary AUC measures. Notably, performance is higher for densities lower than 1% compared to higher densities.

**Fig. S7.**
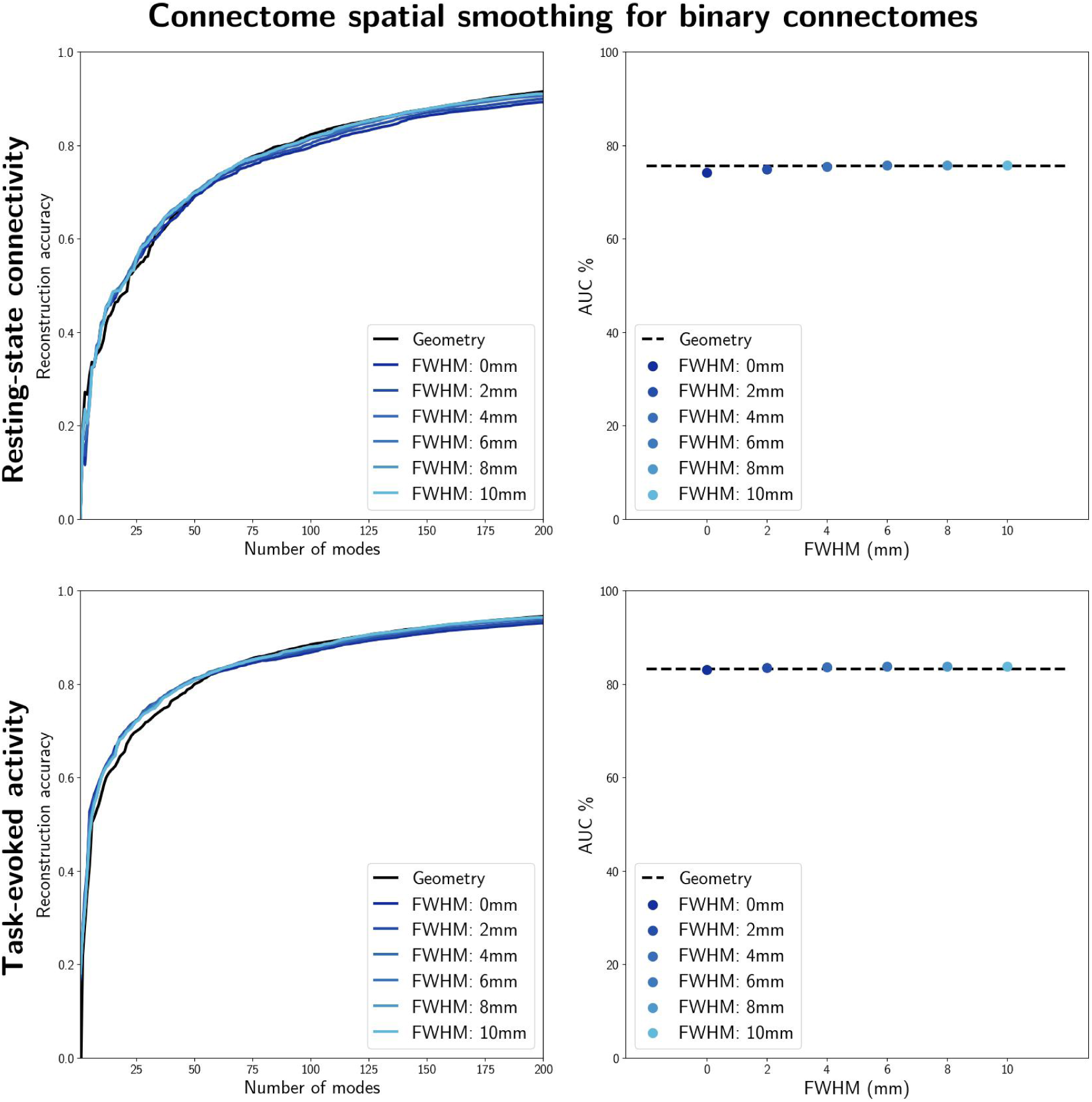
Assessing the impact of connectome spatial smoothing on reconstruction accuracy of binary connectomes. Here, the reconstruction task was systematically repeated for varying smoothing kernels (up to 10mm FWHM) while keeping other pipeline parameters constant; specifically, we utilized the connectomes from our tractography pipeline that better mitigated the gyral bias, performed connectome spatial smoothing, and constructed binary connectomes at 1% density. Line plots (left) depict the reconstruction accuracy as a function of the number of modes for both resting-state connectivity (top) and task-evoked activity (bottom) reconstruction. Scatter plots (right) display the corresponding summary AUC measures. Notably, wider smoothing kernels (greater than 6mmFWHM) yielded slightly improved reconstruction accuracy compared to weaker kernels or no smoothing. Note: The 0mm FWHM case signifies the condition where no smoothing was applied.

**Fig. S8.**
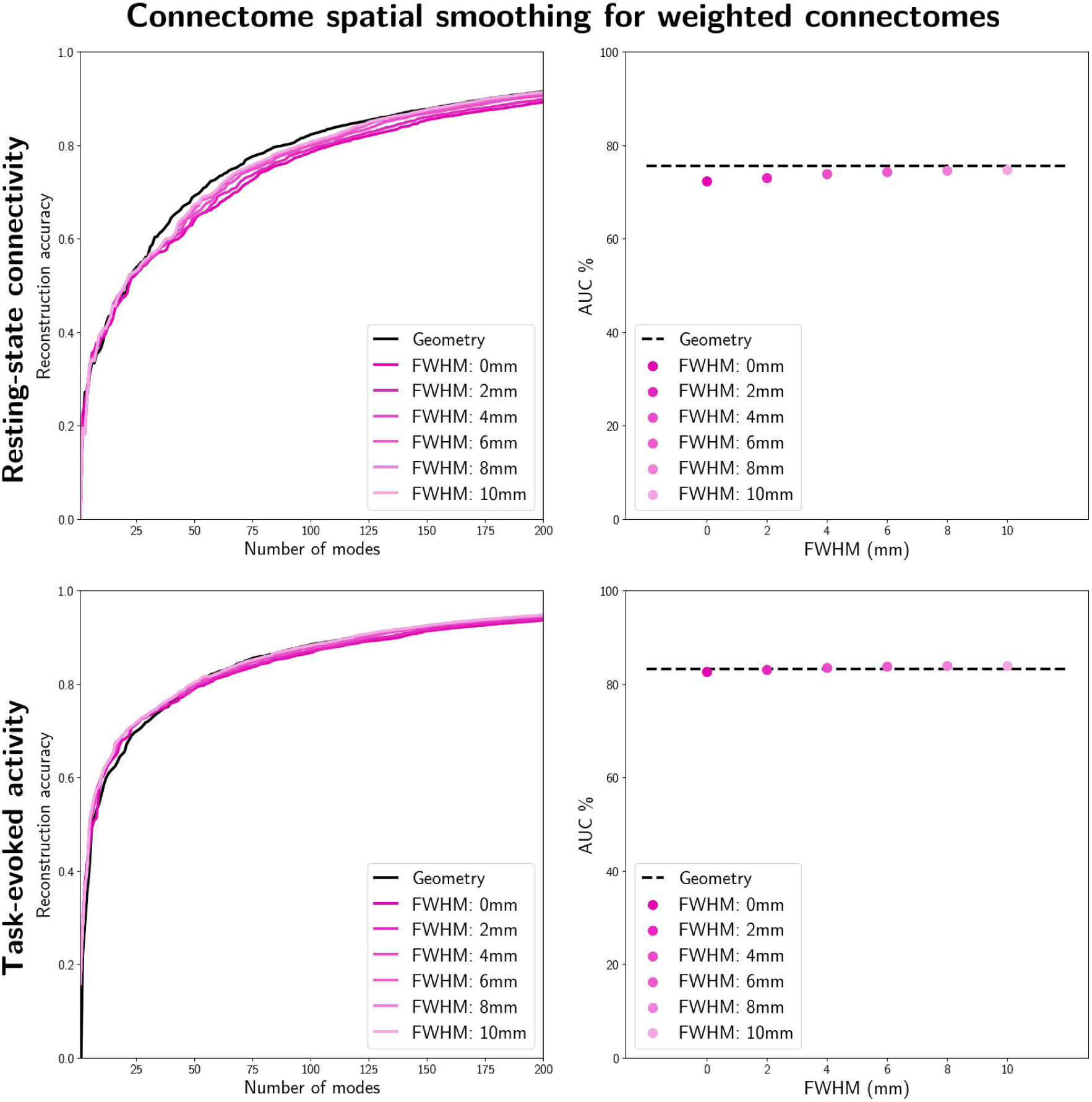
Assessing the impact of connectome spatial smoothing on reconstruction accuracy of weighted connectomes. Here, the reconstruction task was systematically repeated for varying smoothing kernels (up to 10mm FWHM) while keeping other pipeline parameters constant; specifically, we utilized the connectomes from our tractography pipeline that better mitigated the gyral bias, performed connectome spatial smoothing, and constructed a weighted connectome pruned at 10% density. The global-local adjacency combination parameter (ε*_local_*) was fixed at 10^−6^. Line plots (left) depict the reconstruction accuracy as a function of the number of modes for both resting-state connectivity (top) and task-evoked activity (bottom) reconstruction. Scatter plots (right) display the corresponding summary AUC measures. Notably, wider smoothing kernels yielded modestly improved reconstruction accuracy compared to weaker kernels or no smoothing. Note: The 0mm FWHM case signifies the condition where no smoothing was applied.

**Fig. S9.**
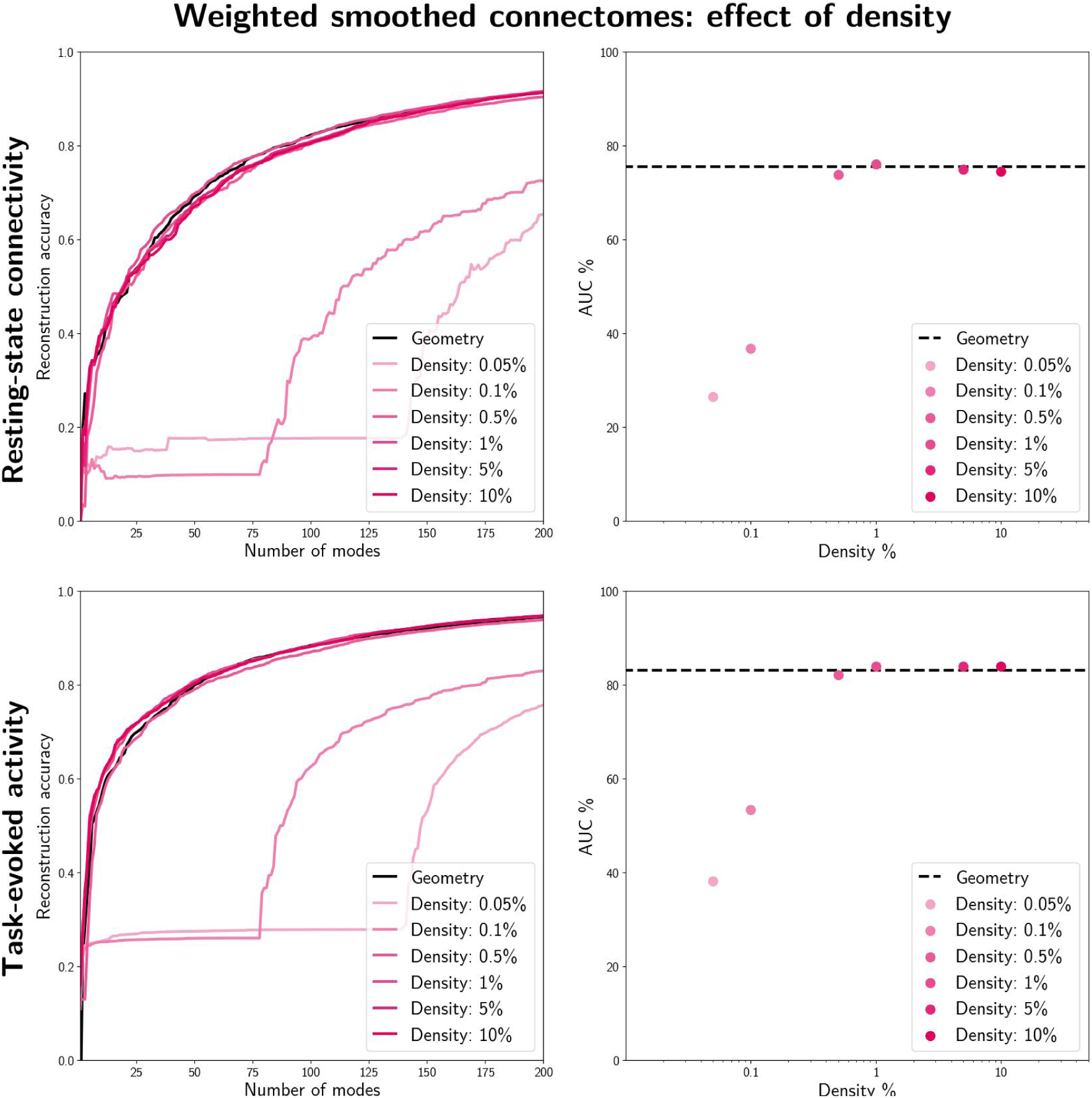
Assessing the impact density on reconstruction accuracy of weighted connectomes. Here, the reconstruction task was systematically repeated for varying density levels (0.05% to 10%) while keeping other pipeline parameters constant; specifically, we utilized the connectomes from our tractography pipeline that better mitigated the gyral bias, performed connectome spatial smoothing (8mm FWHM), and constructed a weighted connectome pruned based on the density criterion. The global-local adjacency combination parameter (ε*_local_*) was fixed at 10^−6^. Line plots (left) depict the reconstruction accuracy as a function of the number of modes for both resting-state connectivity (top) and task-evoked activity (bottom) reconstruction. Scatter plots (right) display the corresponding summary AUC measures. Notably, accuracy is remarkably higher for densities greater than 0.5% compared to lower densities.

**Fig. S10.**
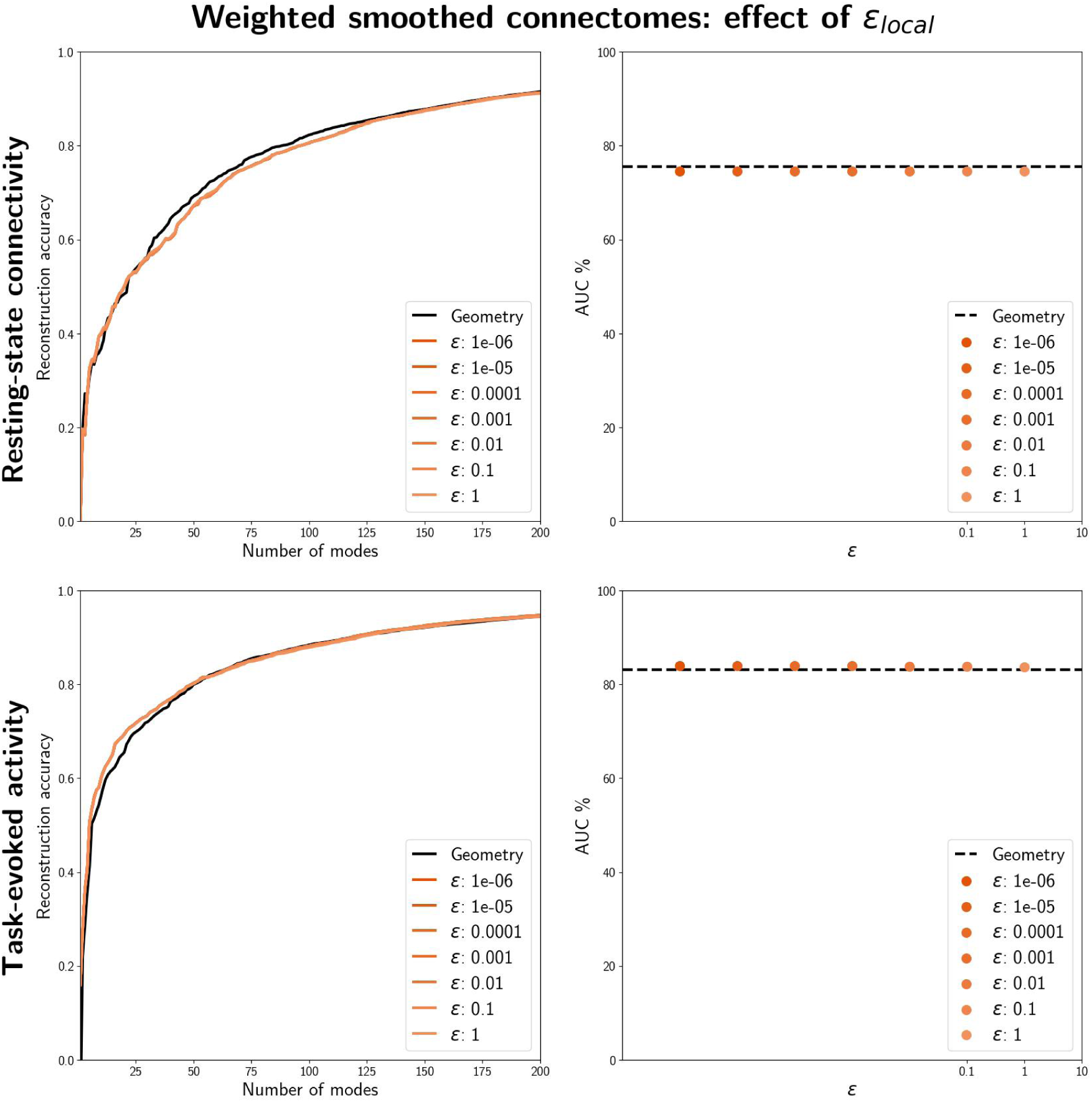
Assessing the impact of the global-local combination parameter (ε*_local_*) on reconstruction accuracy of weighted connectomes. Here, the reconstruction task was systematically repeated for varying choices of ε*_local_* while keeping other pipeline parameters constant; particularly, we utilized the connectomes from our tractography pipeline that better mitigated the gyral bias, performed connectome spatial smoothing (8mm FWHM), and constructed a weighted connectome pruned at 10% density. Line plots (left) depict the reconstruction accuracy as a function of the number of modes for both resting-state connectivity (top) and task-evoked activity (bottom) reconstruction. Scatter plots (right) display the corresponding summary AUC measures. Notably, ε*_local_* seems to have had negligible impact on the reconstruction accuracy.

**Fig. S11.**
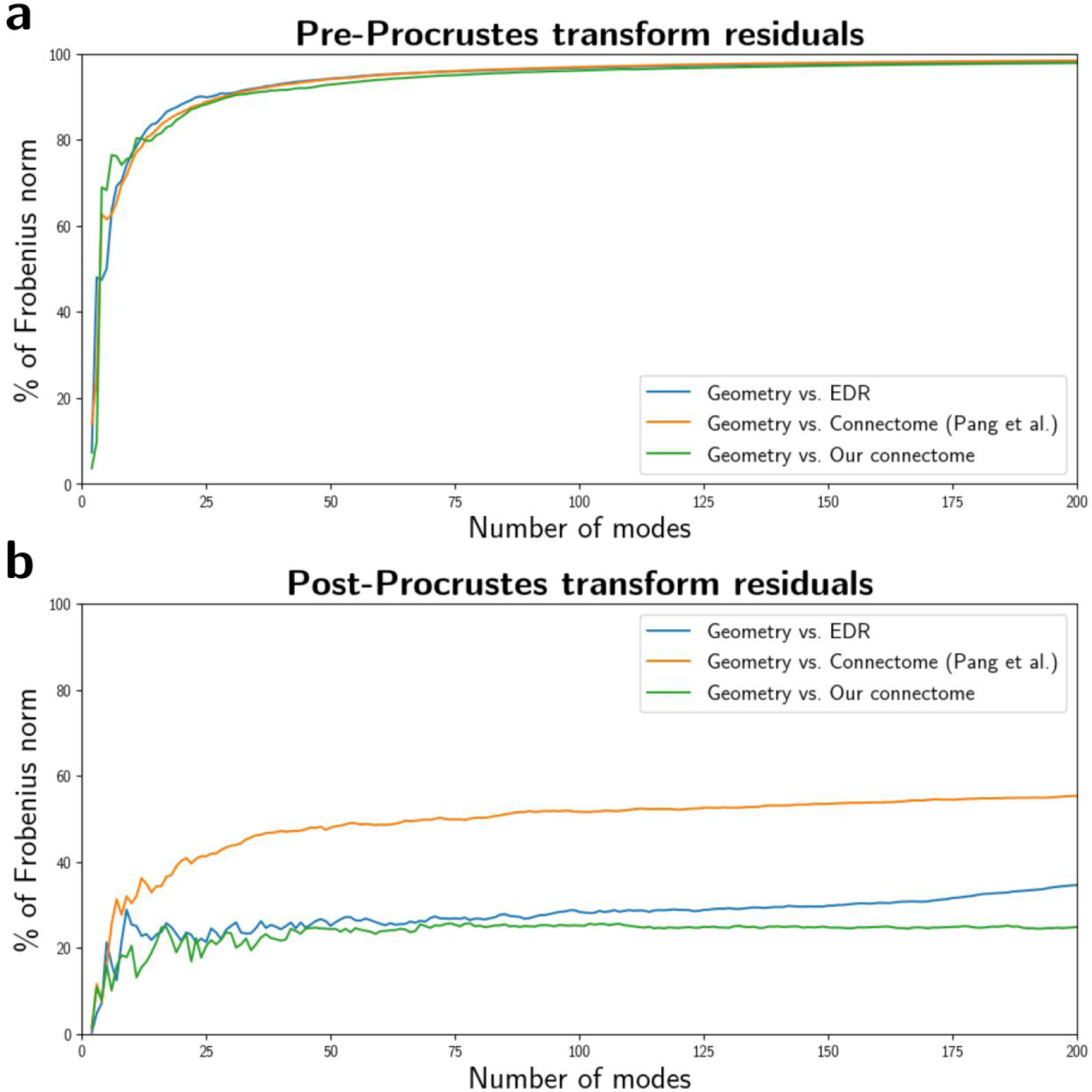
Assessing the distance between pairs of subspaces spanned by different eigenmodes. The x-axis denotes the number of eigenmodes, K, included in the subspace comparison. The y-axis denotes the pairwise eigenmode distances quantified by the Frobenius norm of the off-diagonal values in the K x K cosine similarity matrix. The y-axis is normalized to show the percentage relative to the Frobenius norm of the similarity matrix. (a) Prior to a Procrustes transformation, all pairs of eigenmodes show high distances, i.e. the pairs of subspaces are different due to a lack of alignment between eigenmode pairs. (b) Using Procrustes transformation, pairs of subspaces are optimally aligned. This results in a relative reduction in distance between subspaces. Particularly, distances between pairs of eigenmode subspaces with higher similarity would show a larger relative reduction. These results indicate a higher degree of similarity between EDR/Our connectome eigenmodes to the geometric eigenmodes (in contrast to the lower similarity to Pang et al.’s connectome eigenmodes).

**Fig. S12.**
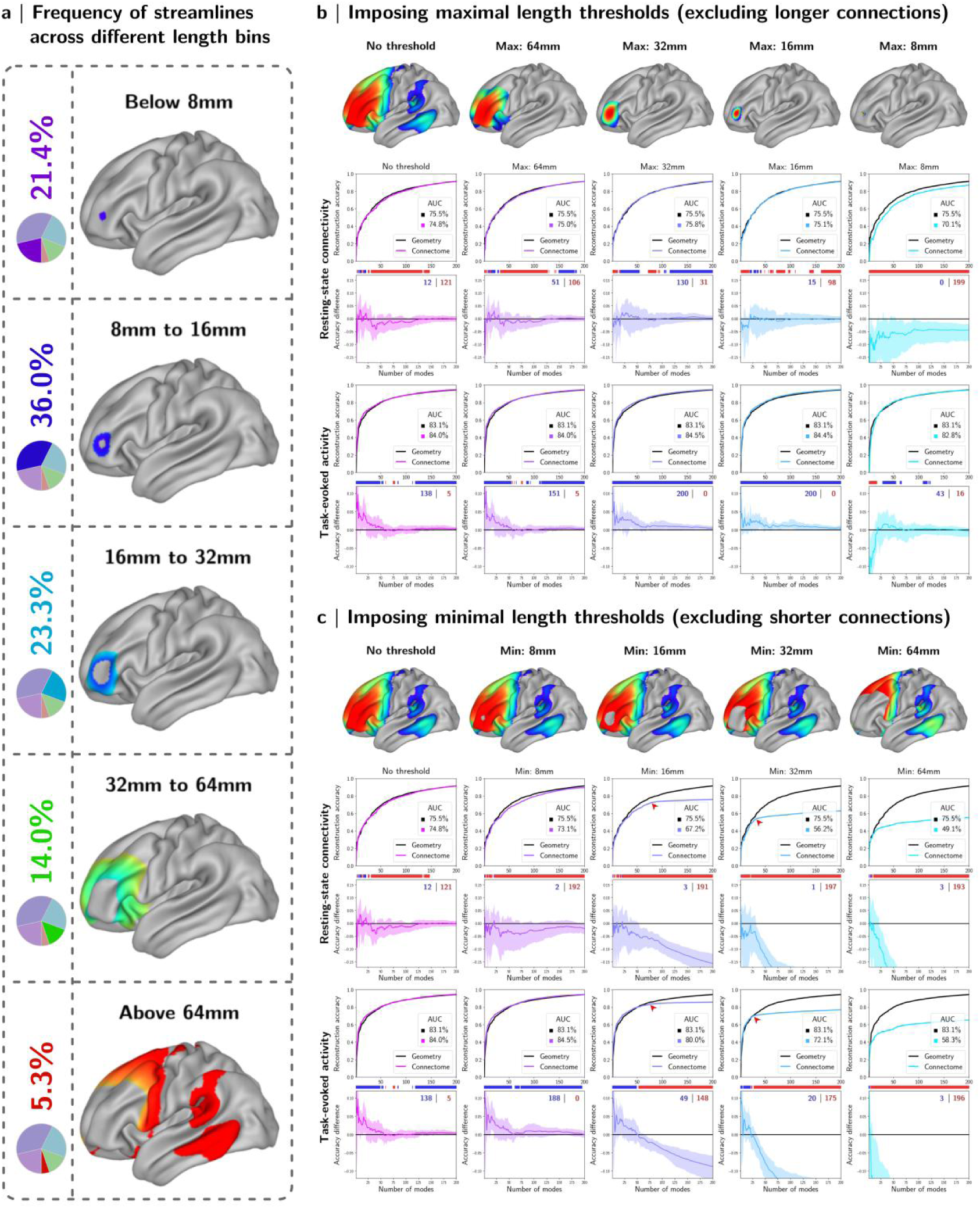
Impact of imposing connection length limits on reconstruction accuracy. (a) Structural connections were grouped based on streamline length. The left panel indicates the frequency of observing streamlines at different length bins along with projections of such connections from an exemplary node. Close to 95% of all streamlines were shorter than 64mm, and more than a fifth of all reconstructed streamlines are shorter than 8mm. (b) The reconstruction accuracy tests were repeated after excluding connections longer than different maximal length thresholds. Imposing maximal length thresholds had relatively modest impacts on reconstruction accuracy, exept for cases with fewer number of modes that were negatively impacted by removal of long connections. (c) The same test was repeated by imposing minimal length thresholds to exclude shorter connections. Exclusion of short connections exerts a more pronounced detrimental impact on reconstruction accuracy, particularly with higher number of modes. This agrees with the intuitive expectation that longe-range connections influence accurate estimation of eigenmodes at longer wavelengths, whereas short-range connections are more influential at constructing eigenmodes at higher spatial frequencies.

**Fig. S13.**
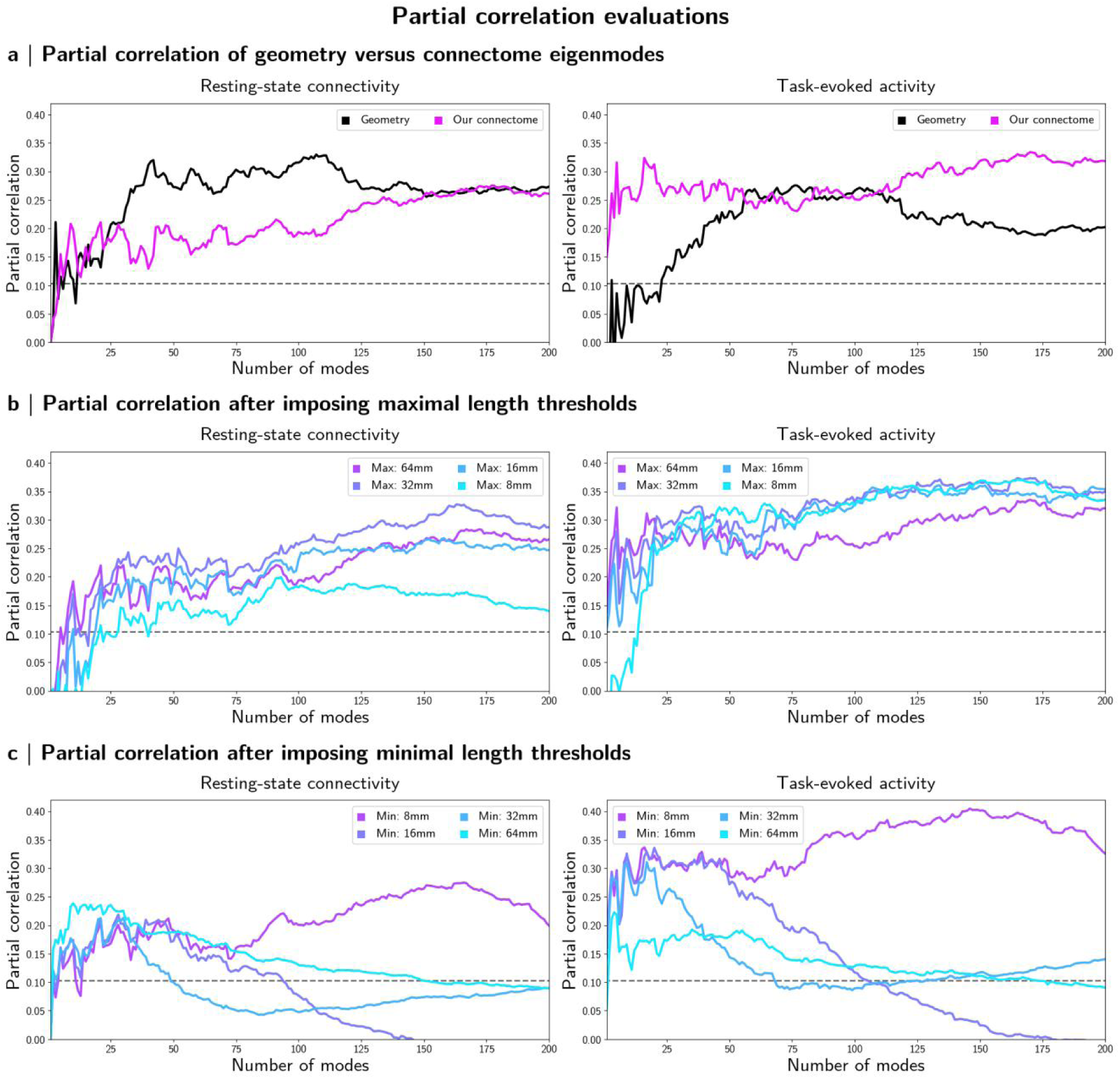
Partial correlation evaluations quantifying the incremental explanatory value of connectome and geometry eigenmodes. Each point represents, for a given basis set & number of modes, the partial correlation between functional activation and the eigenmode-based reconstruction of such, while controlling for those features of the functional activation explained by the alternative basis set using the same number of modes. Dashed gray lines indicate the significance level for a partial correlation test at α = 5%; the null hypothesis of no additional explanatory benefit by the basis set of interest is rejected for points above the line. (a) Comparison between geometric and connectome eigenmodes, using the complete tractography reconstruction in the latter case. The magenta curve represents the scenario where connectome eigenmodes form the basis set of interest while controlling for geometry; conversely, the black curve represents the scenario where geometry eigenmodes form the basis set of interest. (b) The partial correlation assessments after applying maximal length thresholds before constructing connectome eigenmodes. (c) The same test repeated for the case in which a minimal length threshold is instead imposed. Geometry is controlled for all curves in (b) and (c), and the connectome eigenmodes computed after application of the length threshold form the basis set of interest.

